# Three-photon excited fluorescence microscopy enables imaging of blood flow, neural structure and inflammatory response deep into mouse spinal cord *in vivo*

**DOI:** 10.1101/2024.04.04.588110

**Authors:** Yu-Ting Cheng, Kawasi M. Lett, Chris Xu, Chris B. Schaffer

## Abstract

Nonlinear optical microscopy enables non-invasive imaging in scattering samples with cellular resolution. The spinal cord connects the brain with the periphery and governs fundamental behaviors such as locomotion and somatosensation. Because of dense myelination on the dorsal surface, imaging to the spinal grey matter is challenging, even with two-photon microscopy. Here we show that three-photon excited fluorescence (3PEF) microscopy enables multicolor imaging at depths of up to ~550 μm into the mouse spinal cord, *in vivo*. We quantified blood flow across vessel types along the spinal vascular network. We then followed the response of neurites and microglia after occlusion of a surface venule, where we observed depth-dependent structural changes in neurites and interactions of perivascular microglia with vessel branches upstream from the clot. This work establishes that 3PEF imaging enables studies of functional dynamics and cell type interactions in the top 550 μm of the murine spinal cord, *in vivo*.

## Introduction

*In vivo* imaging in the mouse spinal cord has enabled studies of the real-time behaviors and interactions among cells that are associated with homeostatic processes and those linked to disease progression. Examples include imaging neural activity to gain insight into spinal cord sensory neural circuits (Ahanonu et al., 2023; Johannssen & Helmchen, 2010; Ran, Hoon, & Chen, 2016; Sekiguchi et al., 2016), and imaging immune cell invasion and interactions with neurons in various disease and injury mouse models (Caravagna et al., 2018; Davalos et al., 2012; Fenrich et al., 2013; Wu et al., 2022). Such studies capitalize on the capability of nonlinear microscopy to enable micrometer-resolved imaging that can penetrate optically scattering tissue. However, the short optical scattering length of the dorsal spinal cord, due to densely packed myelinated axons, limits the penetration depth for the most commonly used nonlinear contrast mechanism such as two photon excited fluorescence (2PEF) to approximately 150 µm (Cheng, Lett, & Schaffer, 2019; Farrar et al., 2012; Fenrich et al., 2012). This limits studies to only a few superficial regions of the spinal cord, such as the dorsal axonal tracts and restricted grey matter regions outlining the dorsal horn for acute or long-term imaging through window preparations (Farrar & Schaffer, 2014; Lorenzana et al., 2015; Ruschel et al., 2015; Tang et al., 2015; Tedeschi et al., 2019; Wu et al., 2022; Yang et al., 2017). Efforts to visualize cellular functions beyond superficial laminar regions require invasive procedures such as imaging through a chronically implanted microprism (Shekhtmeyster, Carey, et al., 2023) or surgically exposing the spinal cord and angling of the mice to image parts of the ventral horn in an acute preparation (Cartarozzi et al., 2018).

Deeper regions of the spinal cord are anatomically distinct, with different neural functions displaying different vulnerabilities to disease. For example, the dorsal horn neurons accessible with 2PEF microscopy largely encode functions involved in sensation and nociception. Penetration through the dorsal white matter will be necessary to image neural circuits linked with locomotor control. Additionally, the nature of the cellular response to acute insult (e.g. spinal cord injury) or underlying chronic disease states (e.g. multiple sclerosis) varies between laminae within the spinal cord.

Recently, the use of higher order nonlinear excitation of fluorescent markers has enabled deeper imaging into mouse brain. Three-photon excited fluorescence (3PEF) microscopy extends imaging depth over 2PEF microscopy by using a longer excitation wavelength of 1,300-1,700 nm which leads to weaker optical scattering, coupled with the greater confinement of the fluorescence excitation by the higher order nonlinear optical excitation mechanism (Aragon et al., 2022; Chow et al., 2020; Horton et al., 2013; Li et al., 2020; Ouzounov et al., 2017; Wang et al., 2018; Wang et al., 2020; Yildirim et al., 2019). Recently, 3PEF imaging has shown promise for achieving greater imaging depths in the mouse spinal cord (Cheng, Lett, & Schaffer, 2019; Rodriguez et al., 2021).

In this paper, we demonstrate high contrast, micrometer resolution 3PEF imaging to depths of ~550 μm into the spinal cord of living mice with long-term implanted spinal cord imaging chambers. We utilize this imaging technology to map the spinal cord vascular architecture from dorsal venules through capillary beds and to lateral arterioles, as well as to quantify blood flow speed across this microvascular network. We further demonstrate layer-specific responses of neural degeneration after a photothrombotic clot to a surface venule, as well as the time-lapse interactions of perivascular microglia with vessel branches upstream from the occlusion, including microglia migrating toward and invading the vessel lumen.

## Results

### 3PEF using a 1320-nm excitation source enabled imaging to a depth of ~550 µm in the mouse spinal cord *in vivo*

We directly compared the depth penetration of 2PEF (Figure 1A) and 3PEF (Figure 1B) microscopy in the mouse spinal cord by imaging fluorescently labeled blood vessels in anesthetized mice (Movie S1). With 2PEF, imaging with high contrast and resolution was possible in the superficial spinal cord (Figure 1A). With increased imaging depth, background fluorescence (due largely to out-of-focus fluorescence excitation at the sample surface) became brighter (Figure 1C column 1). The signal to background ratio (SBR) became insufficient for reliable imaging at depths below ~150 µm, with the loss of contrast leading to a blurring of feature boundaries. With 3PEF, excitation of out-of-focus background was greatly reduced, enabling high contrast and high resolution imaging of spinal cord capillaries to a depth of ~550 µm (Figure 1C, middle column). Third harmonic generation (THG) from the 1,320-nm pulses enabled visualization of myelinated axons from the white matter (Figure 1C, column 3). Figure 1D shows quantitative comparisons of the image contrast across small spinal cord blood vessels with 2PEF (blue) and 3PEF (magenta) imaging at different depths (Figure 1—figure supplement 1 includes additional, intermediate depths). We found that with 2PEF SBR decayed exponentially with a characteristic attenuation length (CAL) of 50 µm (Figure 1E). In contrast, the 3PEF SBR was relatively constant to a depth of ~400 µm, suggesting background was limited by factors independent of the 1,320-nm laser power, and then decayed with a CAL of 75 µm at greater depth (Figure 1E).

**Figure 1.**
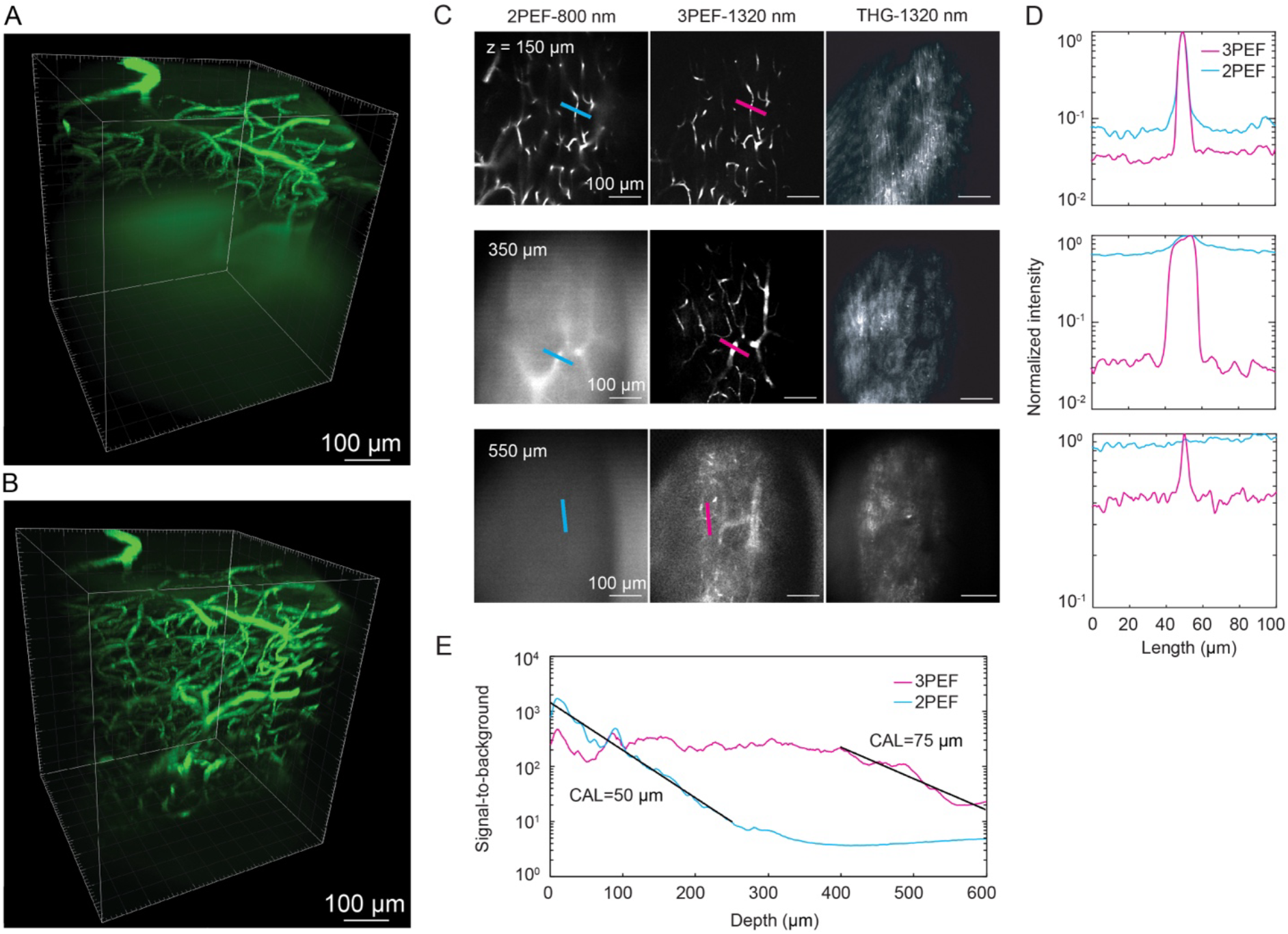
3PEF at 1320-nm excitation wavelength enabled greater imaging depths than 2PEF at 800-nm excitation wavelength in the mouse spinal cord *in vivo.* (A) 2PEF and (B) 3PEF image stacks of fluorescently labeled blood vessels in the spinal cord of a live mouse. (C) 2PEF (left column), 3PEF (middle column), and THG images (right column) at selected depths into the mouse spinal cord. (D) Line profiles across selected capillaries, comparing 3PEF (magenta) and 2PEF (blue) image contrast at different depths. Lines in (C) indicate location of lineouts. (E) SBR for 2PEF and 3PEF imaging of the spinal cord vasculature. In the center region of the frame, the SBR was calculated as the average pixel intensity of the upper 1% to the lower 5%. The line show fits to exponential decays, with the characteristic attenuation length (CAL) of the fit indicated.

### 3PEF imaging allowed mapping of vascular architecture and measurement of blood flow speed from deep arterioles to surface venules in the mouse spinal cord *in vivo*

The dorsal spinal vein (dSV) was apparent along the rostral-caudal axis of the dorsal spinal cord (Figure 2A). High-resolution 3PEF image stacks revealed ascending venules fed by deeper lying capillary beds (Figure 2B), and the dorso-lateral arterioles (dLA) could be visualized at the lateral edges of the spinal cord at depths below 300 µm (Figure 2—figure supplement 1). We began with vessels topologically one branch upstream from the dSV and followed the vascular network deeper into the spinal cord (Figure 2C) upstream into the vascular hierarchy (Figure 2D). We measured vessel diameter (Figure 2E) and quantified blood flow speed by tracking the motion of red blood cells (RBCs) using repetitive line scans along the center of the vessel (Figure 2F) in vessels upstream from the dSV and downstream from the dLA. Flow speed tended to increase, modestly, with increasing vessel diameter in arterioles, and varied little as a function of diameter in venules (Figure 2G). Capillaries, defined as vessels with diameter < 10 µm, had the slowest RBC flow speeds of around 1 mm/s, on average (Figure 2G). Analyzed by topological connectivity, we found capillary flow speeds decreased modestly when moving downstream from arterioles (Figure 2H) and when moving upstream from venules (Figure 2I).

**Figure 2.**
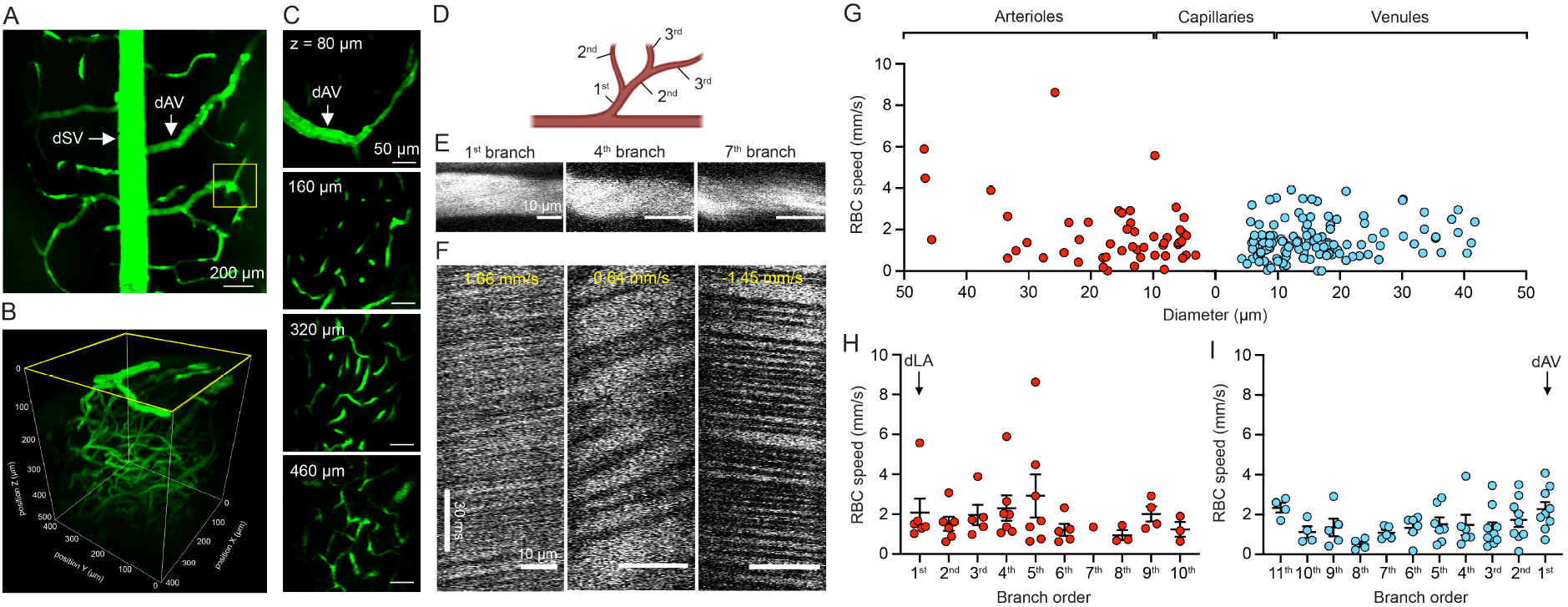
Chronic 3PEF measurement of spinal blood flow from surface venules to deep arterioles *in vivo*. (A) 4x magnification image taken from the dorsal aspect of the mouse spinal cord using 2PEF at 800 nm; spinal cord vasculature was labeled with i.v. injection of FITC-dextran dye (dSV: dorsal spinal vein; dAV: dorsal ascending venule). (B) Volumetric reconstruction of a ~500-μm deep stack of spinal vascular architecture using 3PEF at 1320-nm. (C) Representative images of vascular features at different depths into the spinal cord. (D) Schematic representation defining vascular branch order. (E) Selected vessel segments and (F) corresponding linescan profile from vessels that were one, four, and seven branches upstream from the dAV. (G) RBC speed in arterioles, capillaries (vessel diameter < 10 μm) and venules, expressed as a function of vessel diameter with diameter decreasing in arterioles then increasing in venules from left to right on the graph (n = 166 vessels from 17 mice). Capillaries were assigned to the arteriole or venule side based on their closest topological proximity to an arteriole or venule. (H) RBC speed of arterioles as a function of vascular branch order starting from dLA and going downstream. (I) RBC speed of venules as a function of vascular branch order starting from dAV and going upstream (n = 6 mice).

### A surface venule occlusion led to rapid and progressive neural dieback and inflammatory response across laminar layers in the mouse spinal cord

To demonstrate the application of *in vivo* spinal cord imaging with 3PEF to reveal cellular dynamics in response to injury, we next studied the changes in neurite morphology and the migration of resident microglia in response to the clotting of an ascending venule at the spinal cord surface. Using mice that expressed YFP in dorsal axons, GFP in microglia (Movie S2), and which received retro-orbital injection of red-emitting fluorescent quantum dots to label the vasculature, we took image stacks before, and up to 120 min after photothrombotic clotting of a venule one branch upstream from the dSV (Figures 3A and B; Movie S3). We visualized the cessation of flow in the targeted venule through either 3PEF imaging of fluorescently labeled blood plasma (with RBCs in negative contrast) or from the bright THG signal from the RBCs themselves (Ahn et al., 2020), which revealed static doughnut-shaped RBCs clogged in the vessel lumen after clotting (Cheng, Lett, & Schaffer, 2019) (Figure 3C). Within half an hour of the surface venule occlusion, we observed swelling of axons lateral to the clot location, followed by progressive dieback of these axons (Figure 3D). This axonal degeneration coincided with an overall increase in the density of resident microglia and changes in their morphology at different depths (Figure 3D). Using the THG signal, we also observed a rapid degradation of the myelin sheath around blebbing axons (Figure 3C).

**Figure 3.**
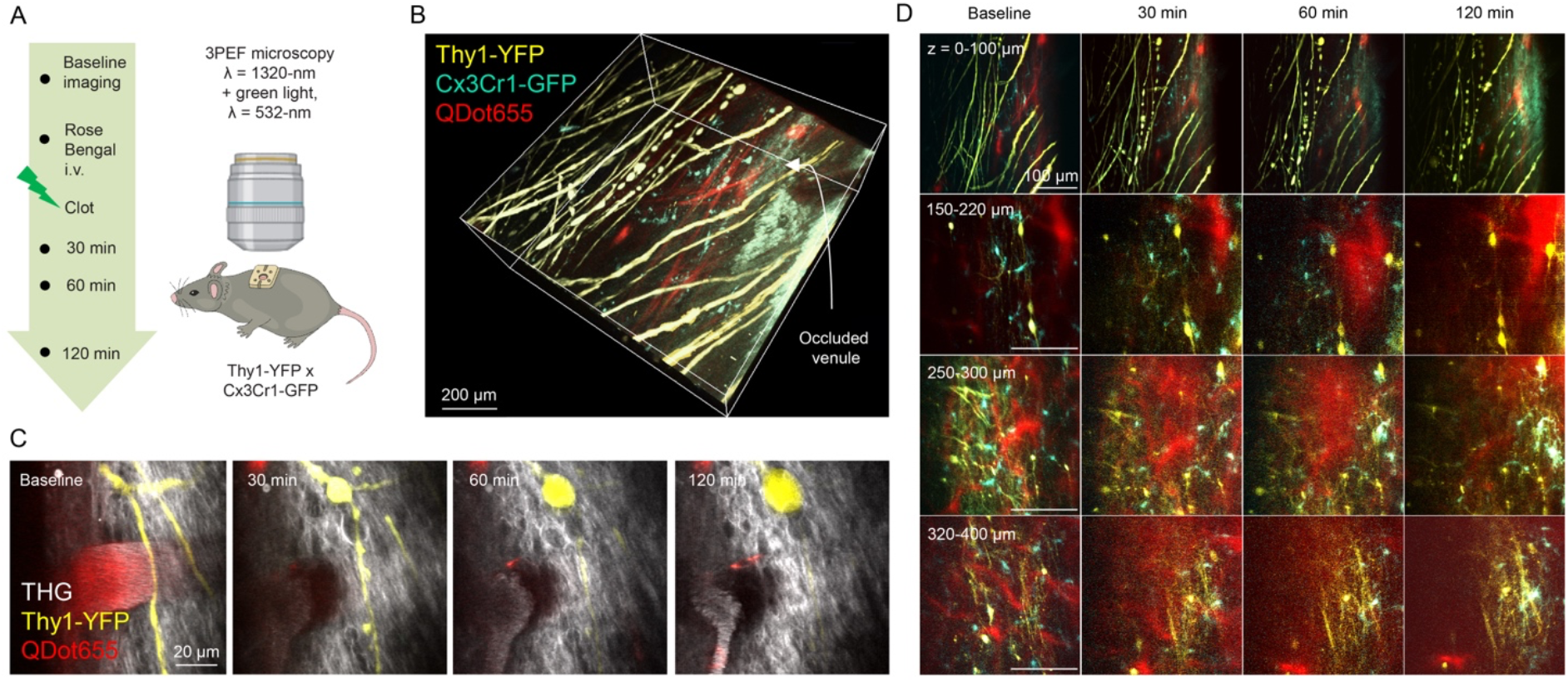
3PEF at 1320-nm enabled multicolor imaging of diverse cellular events in response to a local injury *in vivo*. (A) Experiment timeline and schematics for 3PEF imaging of neural dieback and inflammatory response following a photothrombotic venule occlusion. (B) 3D reconstruction of a z-stack taken from a Thy1-YFP x Cx3Cr1-GFP double transgenic mouse spinal cord; vessels were labeled with i.v. injection of Qtracker^TM^ 655 red-emitting quantum dots. Arrow points to the occluded vessel segment. (C) A cessation of blood flow after a green laser-induced photothrombotic occlusion was followed by rapid axon dieback (yellow, seen via 3PEF) and myelin degeneration (grey, seen via THG) nearby the targeted vessel segment. (D) Time-series of representative images across different laminar depths showing progressive neural dieback, inflammatory response, and capillary disruption after a surface venule occlusion.

### Diverse morphological and behavioral dynamics of neurites and perivascular microglia under acute ischemic conditions

Using a standard system for scoring axon morphology over the course of degeneration (Williams et al., 2014), we observed an irreversible series of morphological changes over 2 hours from intact, swollen, broken, to disappeared (Figure 4A), with a trend toward decreasing intact neurites as a function of time after the occlusion that was more pronounced at greater depth into the spinal cord (Figures 4B, C, D and E). Finally, we evaluated the interaction of microglia with the vessels upstream from the occlusion, motivated by work showing that resident microglia assist in maintaining vascular integrity in ischemic brains (Barkauskas et al., 2013; Fumagalli et al., 2013). We measured microglia dynamics after the occlusion along a continuum of their interaction with blood vessels upstream from the clot, including surface venule branches and small capillaries (Figure 5A). While a proportion of the microglia that were adjacent to vessels remained in a resting state, most migrated toward and extended their dendritic processes toward the vessel, with some progressing to close adhesion, and a smaller subset ultimately invading the vessel lumen, associated with capillary disruption and blood-spinal barrier leakage (Figure 5B). These microglia/vascular interactions tended to be more invasive deeper into the spinal cord laminae after the occlusion as we observed more events associated with perivascular invasion followed by plasma leakage at greater depth (Figures 5C, D, E and F).

**Figure 4.**
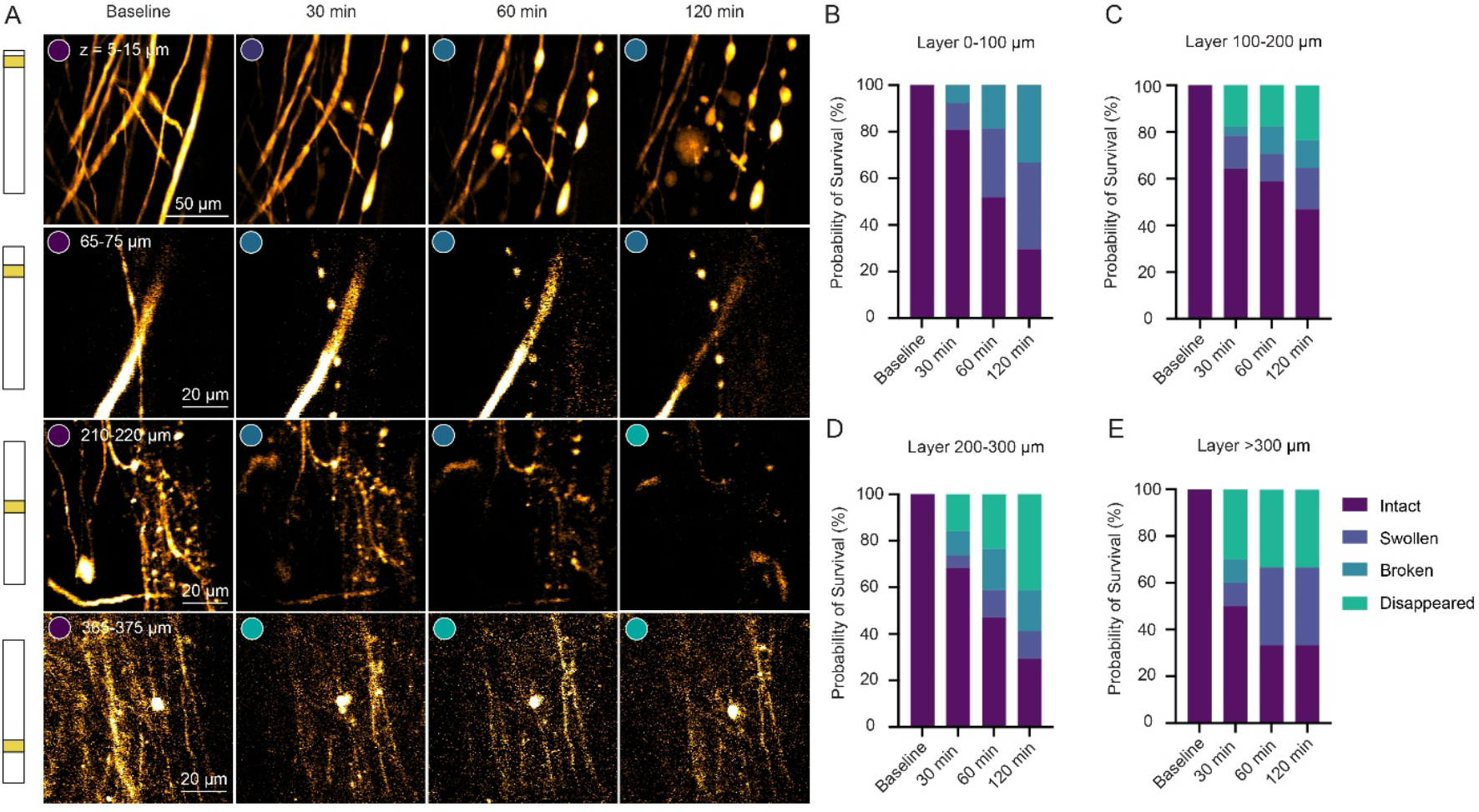
Rapid neurite dieback after a surface venule occlusion. (A) Time-series of representative Thy1-YFP labeled neural structure dynamics from intact, swollen, broken to disappeared across different depths after a photothrombotic stroke. Stacked bar graphs showing the proportion of neural structures that were intact, swollen, broken, or disappeared at different time points from depths of (B) 0-100 μm, (C) 100-200 μm, (D) 200-300 μm, and (E) >300 μm.

**Figure 5.**
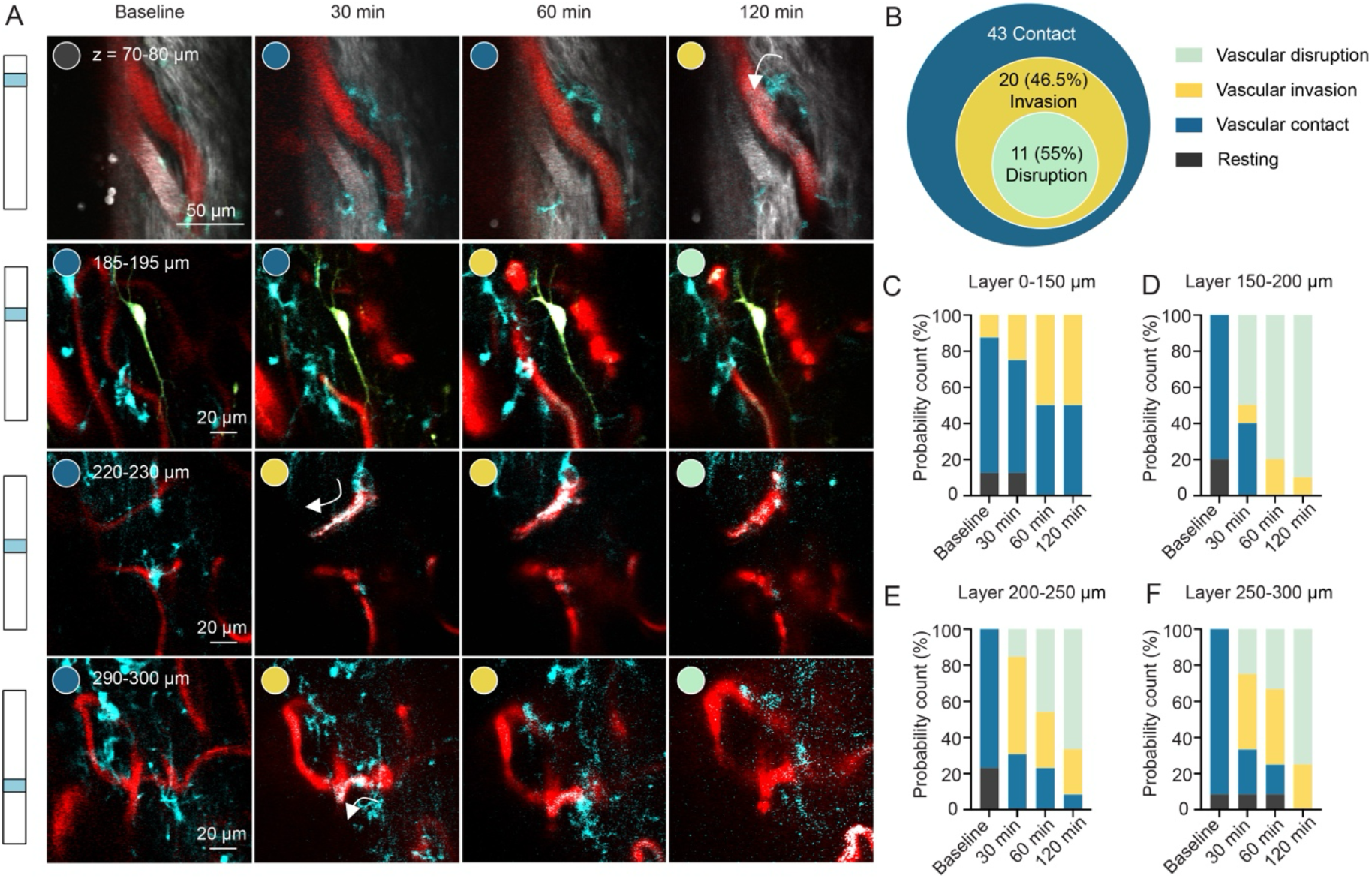
Perivascular Cx3Cr1^+^ cells marched towards vessel lumens, invaded and disrupted small vessels under ischemic conditions. (A) Perivascular Cx3Cr1^+^ cells marched towards vessel walls and extended processes, infiltrating into the vessel lumen (white arrows) and disrupting blood flow after a photothrombotic stroke in an upstream dAV. (B) Summary chart of the behavior of 43 perivascular Cx3Cr1^+^ cells. Perivascular microglia behaviors at depths of (C) 0-150 μm, (D) 150-200 μm, (E) 200-250 μm, (F) 250-300 μm before, 30 min, 60 min, and 120 min after a surface venule occlusion.

## Discussion

The use of optical techniques is especially challenging in the mouse spinal cord due to the highly optically scattering nature of white matter on the dorsal surface (Cheng, Lett, & Schaffer, 2019). Here, we show that 3PEF microscopy overcomes these limitations and enables high contrast and low background 3D imaging of fluorescently labeled structures to depths of ~550 µm into the spinal cord of mice with micrometer spatial resolution and a ~500 µm by ~500 µm field of view. In comparison, 2PEF imaging in the murine spinal cord has been limited to depths shallower than ~150 µm, with rapid degradation of the SBR with imaging depth.

Using 3PEF imaging, we characterized blood flow speed across different classes of microvessels in the spinal cord. We found a modest increase in flow speed with increasing vessel diameter in arterioles (but little speed variation across different diameter venules), contrasting with more pronounced diameter-dependent flow speeds in both arterioles and venules in the mouse cortex (Santisakultarm et al., 2012). Capillary flow speeds averaged around 1-2 mm/s, about the same as have been reported in the cortex (Santisakultarm et al., 2012). With the excitation laser and wavelength used in these studies there is also a strong THG signal from RBCs (Ahn et al., 2020), providing an intrinsic signal that could enable vascular imaging and blood flow speed measurement while “saving” the green emitting fluorophore to label other features. We note, however, that it may be difficult to easily identify small blood vessels with the THG signal in the dorsal white matter, where very strong THG is produced by adjacent myelinated axons.

After photothrombotic occlusion of an ascending venule on the spinal cord surface, we observed progressive degeneration of neural processes in the tissue volume that was drained by the clotted venule, similar to previous findings in the cortex after focal stroke (Shih et al., 2013; Zhang & Murphy, 2007; Zhang et al., 2020). Our data suggest more rapid neural degeneration at greater tissue depths, potentially due to a greater susceptibility of the fine neural processes responding to the ischemic environment in the grey matter as compared to the larger axons on the dorsal surface, or to more severe ischemia at depth after a dAV occlusion. Vessel-adjacent microglia in the capillaries upstream from the occluded venule showed dynamic interactions with the vasculature, migrating toward and even invading capillary segments upstream from the clot, similar to recent findings on behavior of microglia in the cortex after a focal stroke (Barkauskas et al., 2013; Halder & Milner, 2019; Jolivel et al., 2015). The invasion of microglia into the lumen of a vessel was correlated with locally increased leakage of the fluorescently-labeled blood plasma into the parenchyma, although whether this blood-spinal cord barrier compromise is caused by invading microglia or whether it triggers such microglia behavior remains unclear and requires further investigation.

Recent advances in multiphoton microscopy utilizing adaptive optics (AO) to correct tissue-induced aberrations provide improved resolution with deep tissue imaging, and therefore facilitate observing dynamic events that require high SBR and fine spatial resolution (Chen et al., 2021; Liu et al., 2019; Rodriguez & Ji, 2018; Sinefeld et al., 2022; Turcotte et al., 2019). 3PEF imaging with AO was recently shown to enable recording of sensory-evoked calcium responses at the depth of ~300 μm in the mouse spinal cord (Rodriguez et al., 2021). Because of its cylindrical shape and the high refractive index of the lipid in myelin, optical aberrations are expected when imaging into the spinal cord, so AO may provide improved SBR and depth penetration beyond that shown here. The laser powers needed for the deepest penetration into the spinal cord approach the range where tissue injury from residual linear absorption is possible (Wang et al., 2020), although we saw no apparent signs of tissue damage, *in vivo*.

In conclusion, the greater imaging depth afforded by 3PEF imaging opens the door to cell-resolved studies of a spectrum of physiological processes in the spinal cord, as we have demonstrated for measurements of blood flow, neurite degeneration, and inflammatory cell interaction with blood vessels within a context of a focal ischemic injury *in vivo*.

## Materials and Methods

### Transgenic mice

We used 23 mice in total in this study. Wildtype C57BL/6 mice (~2-6 months of age, 20 – 30 g in weight) were used for vasculature imaging and blood flow measurement. For experiments tracking the morphology of neural structures and response of microglia after a photothrombotic occlusion, we used double transgenic mice obtained by crossing Thy-YFPH mice, in which a subset of dorsal spinal cord axons and neurons express yellow fluorescent protein (stock#3782; JAX), with homozygous Cx3Cr1-GFP mice, in which the Cx3Cr1 fractalkine receptor was replaced with GFP leading to fluorescent labeling of resident microglia as well as blood-born monocytes (stock#5582; JAX).

### Spinal imaging window preparation

All animal procedures performed were approved by the Cornell Institutional Animal Care and Use Committee (protocol #: 2015-0029) and were performed under the guidance of the Cornell Center for Animal Resources and Education. To prepare a chronic imaging window for the mouse spinal cord, we followed the window implant procedures previously described (Farrar et al., 2012; Farrar & Schaffer, 2014). Briefly, mice were anesthetized under 1.5% isoflurane in 100% oxygen throughout the entire spinal window implant procedure and were placed on a feedback-controlled heating pad that maintained body temperature at 37°C (50-7053P; Harvard Apparatus). Mice were given i.p. dexamethasone (0.025mg/100g; 07-808-8194, Phoenix Pharm Inc.) and i.p. ketoprofen (0.5 mg/100 g; Zoetis Inc.) to mitigate postoperative inflammation and pain. Atropine sulfate (0.005 mg/100g; 54925-063-10, Med-Pharmex Inc.) was given i.m. to reduce the buildup of mucus in the airways. We injected 0.1 ml of 0.125% (v/v) bupivacaine subcutaneously at the site of skin incision followed by surgically opening the skin above the lower thoracic/upper lumbar spine. Incongruous tissues were gently cleared and removed from the overlying vertebrae, and we then clamped three vertebrae with machined stainless-steel bars. A laminectomy was performed at the T13-L1 vertebral segment using vanna scissors, and small pieces of gel foam along with sterile saline were applied to control bleeding. The dura was left intact. Minimal amounts of Vetbond^TM^ adhesive (1469C, 3M) were carefully applied around the edge of the laminectomy to seal the chamber and further stabilize the attachment of the metal bars. Once the clamping was stabilized and bleeding was controlled, we positioned a top plate with four screws inserted into the metal bars. A small amount of Kwik-Sil^TM^ silicone elastomer (World Precision Instruments) was infused into the well of the chamber and covered with a 5-mm diameter glass coverslip (7229605, Electron Microscopy Sciences). The skin was sealed with Vetbond^TM^ Adhesive, then a layer of C&B MetaBond (Parkell Inc.) was applied at the rostral and caudal side of implant. Animals were allowed to recover for least 10 days, allowing time for post-surgical inflammation to resolve, before any imaging experiments were conducted.

### Excitation source

We used a commercial optical parametric amplifier (OPA) to generate 1,320-nm wavelength pulses for 3PEF and third harmonic generation (THG) imaging (Opera-F, Coherent). The OPA was pumped by a diode-pumped solid-state femtosecond laser (40 μJ/pulse; Monaco, Coherent). The OPA provided an average power of 1.1 W at a 1-MHz repetition rate, with a spectral bandwidth that supports a pulse duration as short as ~50 fs. We adjusted the spacing between an SF11 prism pair, arranged in a standard prism compressor configuration, to maximize the 3PEF signal and thus compensate for the dispersion of the microscope. For 2PEF excitation we used 800-nm wavelength pulses from a commercial Ti:Sapphire laser (Vision II, Coherent).

### Imaging setup

All the images were acquired with a locally built nonlinear microscope. Pulses from the OPA were sent through a computer-controlled rotating waveplate and polarizer to control power, then through a telescope to adjust the beam diameter, and then to scan mirrors to control the scan. Images were acquired using a high numerical aperture microscope objective (25x/1.05-NA, Olympus XLPLN25XWMP2), or with a low magnification objective to take wider field of view images used for orientation (4x/0.28NA, Olympus XLFLUOR). The beam size was set approximately 60% of the back aperture of the 25x objective was filled, which has been shown to be optimal for maximum imaging depth in scattering samples with 3PEF (Wang, Liang, & Qiu, 2015). We could deliver a maximal power of ~230 mW at 1,320 nm through the objective. The emitted florescence and THG was first separated with a 720-nm long pass dichroic (FF01-720/24-25, Semrock), and then split into three channels using two long-pass dichroic filters at 473 nm and 593 nm (Di03-R473 and FF593-Di03, Semrock), and four band-pass filters (center wavelength/bandwidth): 417/60 nm (THG), 494/41 (GFP), 540/80 (YFP, FITC) and 630/92 (Qtarcker655^TM^) (FF01-417/60, FF01-494/41, FF01-540/80, and FF01-630/92, Semrock). Optical signals were collected using GaAsP PMTs (H7422P-40, Hamamatsu). The PMT electronic signals were amplified with a pre-amp with a transimpedance gain of 2 ξ 10^5^ Θ and a 10-MHz bandwidth, and then low-pass filtered with a cut-off of ~500 kHz (about half the pixel clock) before digitization. All images were acquired using ScanImage software (version 3.8, Vidrio Technologies) to control sample movement, laser scanning, power control and signal detection. High-resolution structural images were taken with 1024 x 1024 pixels at 0.8 frames per second. To track RBC flow along individual vessel lumen, we repetitively scanned a bi-directional line across the full frame (1024 pixels) at a line rate of 1.7 kHz for approximately 60 s.

### 3PEF structural imaging

For structural imaging experiments, mice were anesthetized with ~1.5% isoflurane in oxygen-enriched medical air (~40% oxygen) and placed in a custom-built stereotaxic apparatus with a feedback-controlled heating pad set at 37°C (50-7053P; Harvard Apparatus). To map the vascular architecture in the spinal cord, the blood plasma was labeled with a 100-μL retro-orbital injection of 5% W/V FITC-dextran dye (MW=70,000 kDa; Invitrogen), or Qtracker^TM^ 655 (20 μL in 100 μL saline; Invitrogen). Respiration was carefully monitored during the imaging session and the isoflurane level adjusted to maintain a steady breathing rate of ~1Hz. We gave the mouse an hourly dose of i.p. 5% W/V glucose in saline (1mL/100g) to maintain hydration and i.m. atropine (0.001mg/100g) to prevent lung secretions. Image stacks were taken through the 25x water-immersion objective using a 1,320-nm excitation wavelength. We acquired 1-μm step size image stacks beginning with a laser power of ~10 mW at the surface of the spinal cord tissue. The beginning of the z-stack was determined by the depth at which the first vessel that came into focus in the FOV. We manually increased the power as imaging depth increased to maintain the 3PEF signal until 230 mW was used at the maximum imaging depth of ~550 μm. The SBR was calculated as the ratio between the average value of the brightest 1% of pixels (signal) divided by the average value of the 5% dimmest pixels (background) from a region at the center of the field of view, at each depth. We fit the SBR to an exponential decay curve to determine the CAL (Wang et al., 2020).

### Characterization of blood flow speed from surface venules to deep arterioles

We measured blood flow speed beginning with ascending venule vessel segments that branch from the prominent dorsal spinal vein (dSV), and following the vascular tree, branch-by-branch deeper into the tissue – and upstream along the vascular topology – until image contrast was lost. For each vessel segment we measured, vessel diameter was first determined by maximum intensity projection of a small z-stack, and manual tracing of the vessel boundaries. For measuring the blood flow speed, we used the line-scan approach described previously (Farrar et al., 2015; Santisakultarm et al., 2012). Briefly, the scan pattern was adjusted to repeatedly scan a line down the center of the vessel segment for ~1 min, building a two-dimensional space-time image. Since the injected dye only labels the blood plasma, the movement of individual RBCs produces diagonal dark lines in the space-time image, with a slope inversely proportional to the blood flow speed.

### Photothrombotic stroke model with rose bengal

A 532-nm continuous wave laser (Compass 215M, Coherent) was routed into the 3PEF imaging setup and focused through the microscope objective in the same plane and at the center of the imaging field. Mice were retro-orbitally injected with 50 µL of 10 mg/mL rose bengal in saline. We targeted ascending venules, one branch upstream from the dSV. Once the target vessel was centered in the imaging field, it was irradiated with the green laser at ~5-mW incoming power and for repeat bouts of 10-30 s over 2-3 min, with intermittent imaging to assess clot formation. This continued until a clear clot formed in the targeted venule and there were no signs of blood flow in the target vessel (Schaffer et al., 2006; Zhang et al., 2020).

### Evaluation of neural damage and microglia reactions to spinal cord venule occlusion

We produced photothrombotic occlusions in spinal cord ascending venules in mice expressing YFP in a subset of Thy1-expressing neurons and GFP in Cx3Cr1-expressing resident microglia. We followed the upstream vascular tree from the target vessel and took repeated z-stacks in the region of tissue which was drained by the target venule. These images were taken before, and at 30, 60, and 120 min after the vessel occlusion. In these image stacks, we first identified neural process segments (~70-80 neurites from each mouse, n = 3 mice), including both axons and dendrites, to follow over all imaging time points. We scored the morphology of these segments as intact, swollen, broken, or disappeared, following a previously described scoring system (Rosidi et al., 2011; Williams et al., 2014). Swollen neurites have an expanded diameter without complete segmentation, while broken neurites are defined as having an evidently beaded structure. We also identified microglia that were proximal to vascular branches upstream from the clotted vessel. We followed the migration and subsequent interaction of these microglia with the nearby vessels, and scored this interaction as contacting (extending processes to contact the vessel), invading the vessel (microglia invade and adhere inside the vessel lumen), or disrupting the vessel (evident from fluorescently labeled blood plasma leaking into the parenchyma).

## Funding

National Science Foundation NeuroNex program DBI-1707312 (CBS)

New York State Spinal Cord Injury Research Board C32094GG (CBS) C32630GG (YTC)

National Institutes of Health NS096669 (CBS) Craig H. Neilsen Foundation 296332 (CBS)

Mong Family Foundation, Cornell Neurotech program (YTC)

## Author contributions

Conceptualization: YTC, CX, CBS

Investigation: YTC and CBS designed the research; KLM performed experiments shown in Figure 1; YTC performed experiments shown in Figures 2 - 5.

Supervision: CBS

Writing—original draft: YTC, KML, CBS

Writing—review & editing: YTC, KML, CX, CBS

## Data and materials availability

All data needed to evaluate the conclusions in the paper are present in the paper and/or the Supplementary Materials. Any raw data related to this paper may be requested from the authors.

Figure 2D and Figure 3A were created with BioRender.com.

## Supplementary Materials

**Figure 1—figure supplement 1.**
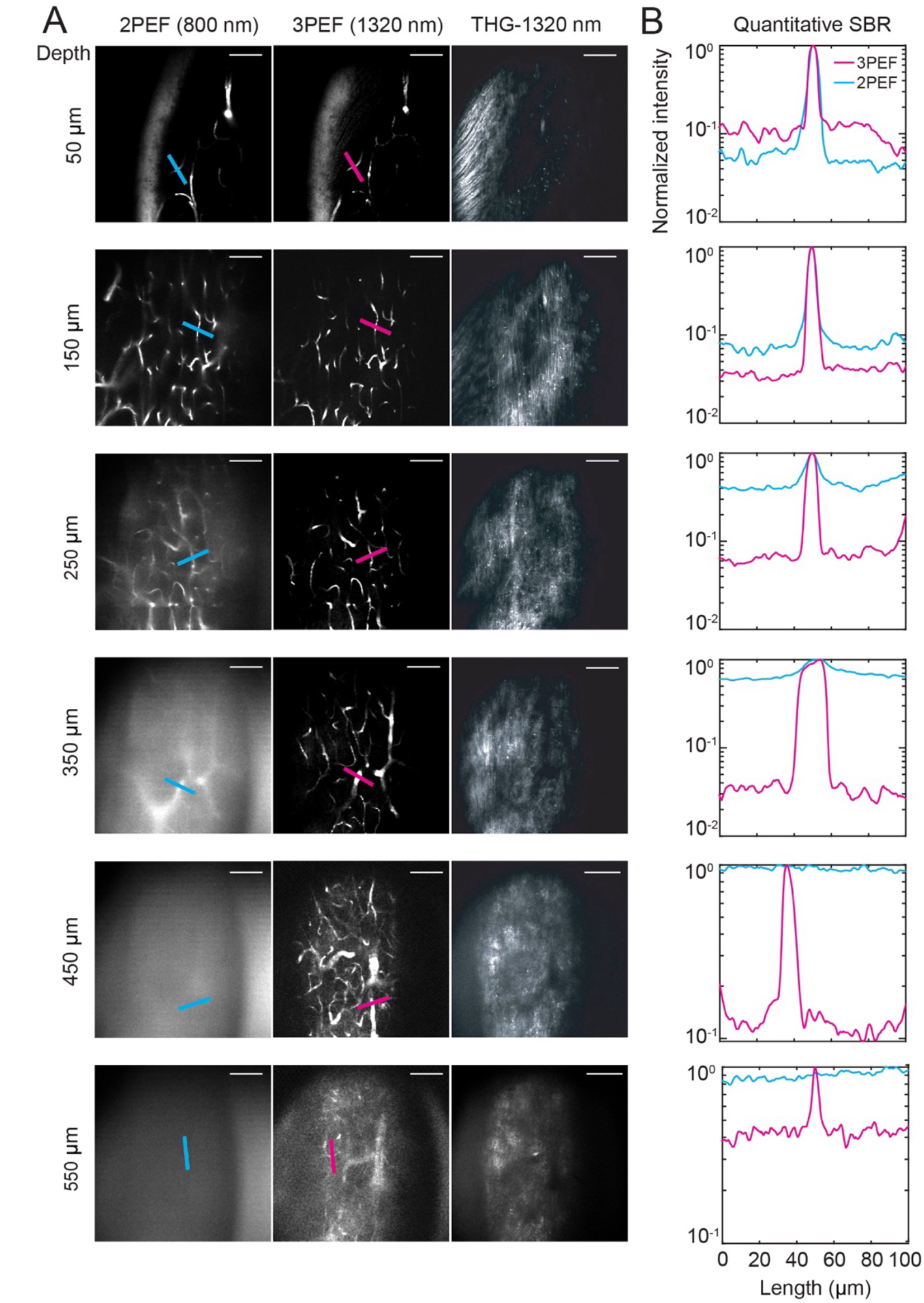
Representative images of fluorescein-labeled vasculature stack under 2PEF at 800-nm versus 3PEF at 1320-nm excitation source. (A) 2PEF (left column), 3PEF (middle column), and THG images (right column) at selected depths into the mouse spinal cord. Scale bar: 100 μm. (B) Line profiles across selected capillaries, comparing 3PEF (magenta) and 2PEF (blue) image contrast at different depths. Lines in (A) indicate location of lineouts. These data include the three depths shown in Figs. 1C and D and add three additional depths. Scale bar: 100 μm.

**Figure 2—figure supplement 1.**
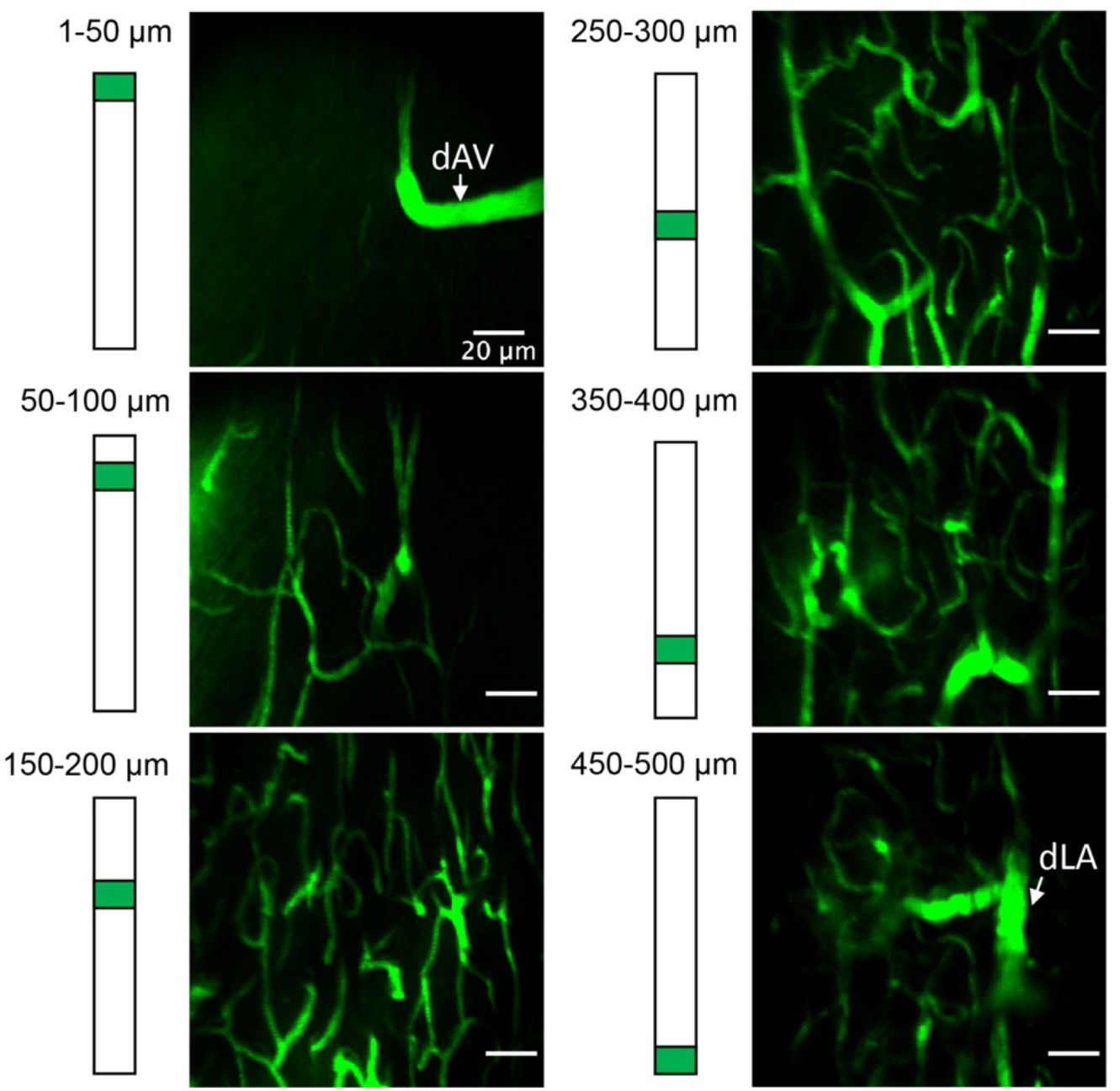
Representative images demonstrating spinal cord vasculature, including dAV at the surface and dLA at depth. Selected images from a continuous vasculature stack indicate the dAV at the top 50 μm followed by a dense capillary network spanning across the spinal cord tissue column, which is fed by dLA that runs along the rostral-caudal axis of the mouse spinal cord.

**Movie S1. Images stacks showing spinal cord vasculature taken with 2PEF (left) vs 3PEF (right).** Stacks correspond to the image data in Figure 1.

**Movie S2. Three-dimensional rendering of 3PEF image stack from Thy1-YFP x Cx3Cr1-GFP mouse showing axons (YFP, shown in green) and microglia (GFP, shown in magenta).** Image stack corresponds to data shown in Figure 3B, taken at baseline, before the dAV occlusion.

**Movie S3. 3PEF image stacks of blood vessels (Qtracker^TM^ 655, shown in red), axons (YFP, shown in yellow), and myelin (THG, shown in greyscale) taken at baseline, and at 30 min, 60 min, and 120 min after the occlusion of a dorsal ascending venule.** Image stacks correspond to the occlusion site shown in Figure 3C.

